# INDOCYANINE GREEN IMAGING BY VITAL TISSUE SLICES SCANNING ALLOWS FOR THE ISOLATION OF INTACT LIVER MICRO-METASTASIS FOR SINGLE-CELL ANALYSIS

**DOI:** 10.1101/2023.06.21.545927

**Authors:** Niccolò Roda, Massimo Stendardo, Valentina Gambino, Giuseppe Persico, Andrea Cossa, Simona Rodighiero, Chiara Priami, Nina Kaludercic, Pier Giuseppe Pelicci, Marco Giorgio

## Abstract

Metastasis represents the deadliest outcome in cancer, leading to the vast majority of cancer-related deaths. Understanding the progression from micro-to macro-metastasis might improve future therapeutic strategies aimed at blocking metastatic disease. However, the difficulty of investigating vital, clinically undetectable, micro-metastases hindered our capacity to unravel phenotypic determinants of micro-metastases. In this work, we leveraged indocyanine green (ICG) dye to detect small sized liver micro-metastases across several cancer models. We exploited a method for infrared fluorescence scanning of fresh tissue and coring of cancer micro-metastases and succeeded in processing them for single-cell RNA sequencing. Our analysis revealed that distinct liver micro-metastases upregulate both shared and specific genes that can successfully predict breast cancer patient prognosis. Moreover, the ontology classification of these genes allowed the validation of several pathways, namely interferon response, extracellular matrix remodeling, and antioxidant response in metastatic progression. Ultimately, we showed that ICG can be successfully used to quantify breast cancer micro- and macro-metastases to lungs, which we showed to be abrogated through inhibition of H_2_O_2_-producing enzyme monoamine oxidase. Therefore, the ICG approach allowed us to identify not only determinant of breast cancer metastatization, but also to assess the therapeutic efficacy of targeting these genes which can be further investigated in clinic.

## INTRODUCTION

Cancer-related mortality is nearly exclusively due to metastatic disease (**1**), for which systemic treatment is rarely □ if ever □ curative (**2**). In metastatization, primary tumor cells invade the surrounding tissue, intravasate and survive in the bloodstream, extravasate in distant organs where they progress from micro-to macro-metastases (**3**). In this respect, micro-metastases are defined as small deposits of cancer cells in secondary organs, invisible to any imaging tests and only visible by microscopy. Traditionally, metastasis spreading has been envisaged as a late process requiring tumor masses to grow up to a minimal size before releasing cells in the circulation. However, several studies rather suggest that metastasis spreading is an extremely early event, with tumoral cells disseminating from the pre-malignant phase of tumor development (**4**). Coherently, in most cases, metastases appear after the surgical resection of an apparently localized disease, following the growth of clinically undetectable micro-metastases already present at diagnosis (**5**). In this scenario, the progression from micro-to macro-metastasis represents the most important clinical window to improve patient prognosis. Therefore, development of therapeutic strategies aimed at preventing the progression from micro-to macro-metastasis is still an unmet clinical need, mostly because of the insufficient understanding of micro-metastasis biology.

So far, functional studies on micro-metastasis have been performed on macroscopic lesions (1-2 mm size in the best cases), due to the limitations of *in vivo* imaging (**6**). In xenograft models the use of fluorescent labelled tumor cells shed light only on the modality of metastatic dissemination and colonization (**7**). However, in clinical settings, micro-metastasis cannot be investigated below 0.4-0.5 mm size, a resolution achieved by using tissue processing and embedding procedures that kill cells (**8**).

Indocyanine green (ICG) is a 775 Da water-soluble compound displaying fluorescent properties in the near infrared region (IR) (**9**). In clinics, ICG was successfully exploited as vital dye for angiography (**10**) and fluorescence navigation surgery (**11**). Currently, intravenous ICG administration represents the most common tracing strategy for tumor resection (**12**), particularly for liver primary and metastatic lesions (**13**), gastric primary and metastatic lesions (**14**), breast (**15**) and brain (**16**) tumors. Ultimately, ICG was adopted to stain sentinel lymph node in different cancers (**17**).

In pre-clinical settings, ICG has been widely used in oncology studies aimed at investigate *in situ* and metastatic osteosarcoma (**18**) and at detecting colorectal cancer macro-metastases (**19**). However, no previous work leveraged ICG to study micro-metastasis biology.

In this work, we revealed that ICG can successfully stain liver micro-metastasis deriving from different human cancer models and from the spontaneous breast murine model MMTV-WNT-1. In addition, we leveraged ICG imaging to identify, collect, and process breast cancer (BC) liver micro-metastases in different BC models (i.e., orthotopic and intravenous injection of the BC cell line MDA-MB-231, and MMTV-WNT-1 model). Collected cells subsequently underwent single-cell RNA sequencing (scRNA-seq), which allowed us to unravel the transcriptional profile of BC micro-metastases and to investigate the inter-micro-metastasis transcriptional heterogeneity. Ultimately, we showed that ICG can be further exploited to assess the effect of the monoamine oxidase B (MAO-B) inhibitor safinamide (**20**) pre-treatment on BC lung micro-metastases. Therefore, our work revealed a previously unknown application of ICG in the oncology field and identified previously undescribed molecular determinants of BC micro-metastases.

## MATERIALS AND METHODS

### Data availability statement

All the data that support the figures and the other findings are available from the authors upon request.

### Cell lines

All cells were cultured in adhesion in a 20% O_2_, 5% CO_2_ incubator at 37°C. HEK293T and MDA-MB-231 were purchased from the American Type culture collection (ATCC) and cultured in DMEM (EuroClone), supplemented with 10% South American fetal bovine serum (FBS; Microtech), 2 mM L-glutamine (EuroClone), and 100U/ml penicillin-streptomycin (Thermo Fisher Scientific). All cell lines were tested for mycoplasma contamination routinely. All cell lines were split once they reached ∼80% confluence and cultured *in vitro* for no more than 10 passages after thawing. U11 CRC PDX was kindly provided by prof. Ruggero de Maria (Universita’ Cattolica del Sacro Cuore, Rome) and cultured as the cell lines.

To test the effect of safinamide administration on BC metastasis burden in, 1×10^6^ MDA-MB-231 were plated in 10 cm dishes and treated with safinamide (10μg/ml) for 72 hours.

### Mice

Female NOD/SCID Il2-Rγ null (NSG) and FVB MMTV-WNT1 mice were bred in house and housed under specific pathogen-free conditions at 22 ± 2°C with 55 ± 10% relative humidity and with 12 hours day/light cycles in mouse facilities at the European Institute of Oncology–Italian Foundation for Cancer Research (FIRC) Institute of Molecular Oncology (IEO–IFOM, Milan, Italy) campus. *In vivo* studies were performed after approval from our fully authorized animal facility and our institutional welfare committee and notification of the experiments to the Ministry of Health (as required by the Italian Law (D.L.vo 26/14 and following amendments); IACUCs Numbers: 833/2018, 762/15, and 679/20), in accordance with EU directive 2010/63.

### Viral production

Lentiviruses were packaged in HEK293T cells using pMDLg/RRE, pRSV-REV, and pMD2.G third generation packaging and envelope plasmids. The Tet-Off-H2B-GFP lentiviral vector (Falkowska-Hansen) allows the constitutive expression of the fusion protein histone H2B-GFP under the control of a tetracycline-responsive promoter element (TRE), which is in turn regulated by a tetracycline-controlled transactivator protein (tTA). Tet-Off-H2B-GFP lentiviral vectors were transfected through calcium phosphate transfection. Briefly, 10μg of transfer plasmids were mixed with 4μg of pMD2.G, 3μg of pRSV-REV, and 4μg of pMDLg/pRRE packaging vectors per each 10cm dish of HEK293T. CaCl_2_ (Sigma-Aldrich) was added to the vectors mix to a final concentration of 0.5 M. The mix was added dropwise to 500 μl of 2X HBS (HEPES buffered saline (Thermo Fisher Scientific): 250 mM HEPES pH 7.0, 250 mM NaCl and 150 mM Na_2_HPO_4_), constantly bubbling. Chloroquine (Sigma-Aldrich) was added to HEK293T medium to the final concentration of 100μM to increase transfection efficiency. After 15 minutes of incubation at room temperature, the suspension was added dropwise to the plates and immediately mixed with the culture medium by gentle swirling. Viral supernatants were collected 2-4 d post-transfection and filtered through 0.45 mm filters. Filtered supernatants were concentrated by ultracentrifugation at 22,000 rpm for 2 hours at 4°C with OptimaTM L-90K Ultracentrifuge (Beckman Coulter) and stored at -80°C (never refrozen).

### *In vivo* experiments

In the orthotopic transplantation setting, per each mouse, 200,000 breast cancer MDA-MB-231 cells were resuspended in 30 μl of PBS (Thermo Fisher Scientific) and growth-factor-reduced Matrigel (Corning) in a ratio of 1:1. Cells were injected in the ninth mammary gland of 8–12-week-old female NSG mice without prior clearing of the fat pad. Mice were sacrificed at specific time points as discussed in the Results section to investigate liver and lung metastases. The metastatic burden (Fig. 2D and 4B) was computed as the fraction between the total area occupied by metastatic cells in liver (or lungs) and the total area of the liver (or lung) across all sections. This value was normalized on the value at the first investigated time point to obtain the increase in metastatic burden. In the intra-venous setting, per each mouse, 200,000 MDA-MB-231 were resuspended in 200 μl of PBS (Thermo Fisher Scientific) and injected in the caudal vein of 8–12-week-old female NSG mice. Mice were sacrificed 28 days after injection to inspect either liver or lung metastases. In both experimental settings, MDA-MB-231 were first transduced with a lentiviral H2B-GFP reporter construct to allow for their identification *in vivo*. Upon infection, cells expressing high levels of GFP were isolated from dissociated tissues via FACS using a BD FACSAria II.

**Fig. 1:**
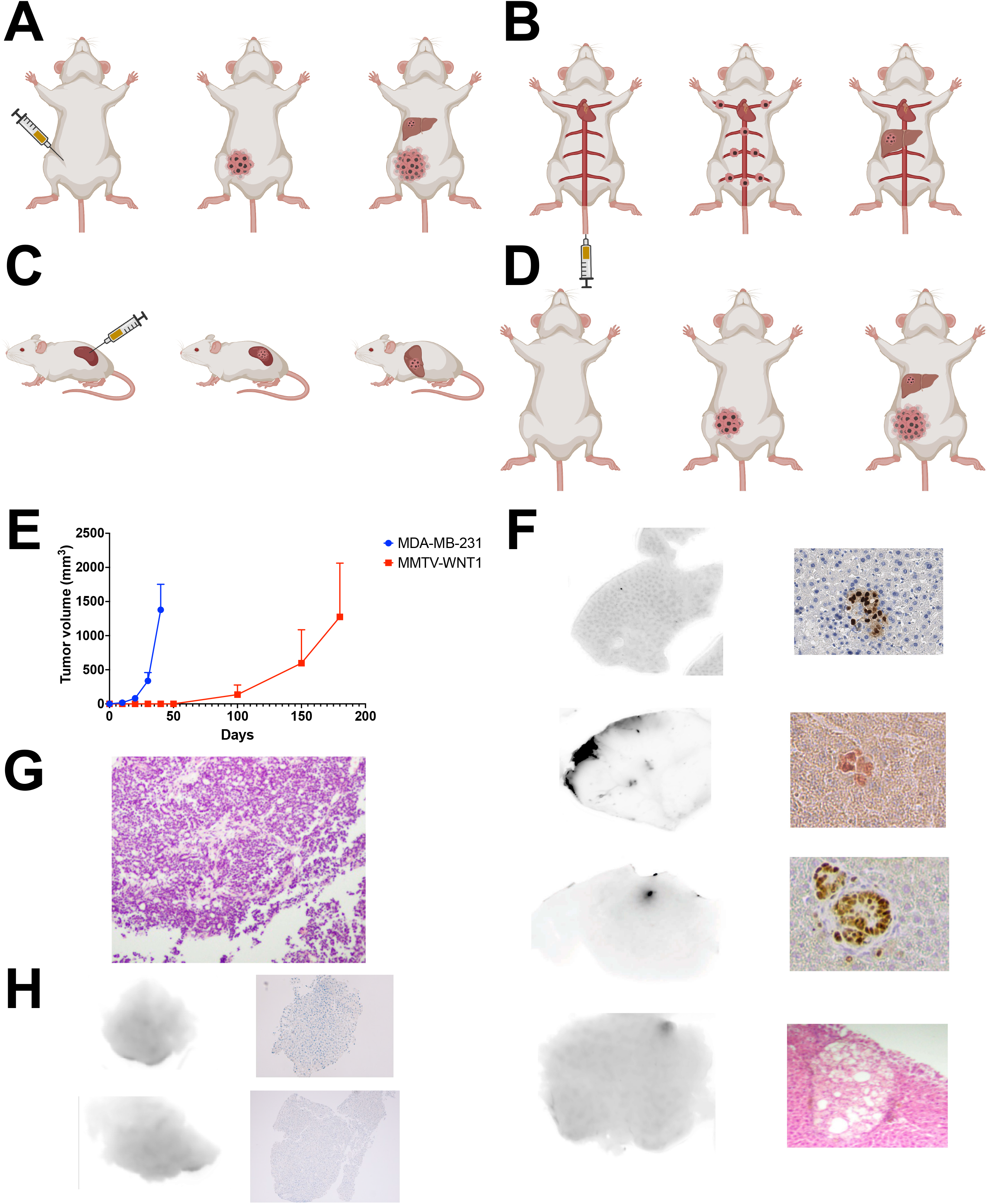
ICG staining allows for visualization of liver micro-metastases. **(A)** Schematic representation of the orthotopic model of BC micro-metastases. **(B)** Schematic representation of the intra-venous model of BC micro-metastases. **(C)** Schematic representation of the intra-splenic model of CRC micro-metastases. **(D)** Schematic representation of the spontaneous model of MMTV-WNT1 BC micro-metastases. **(E)** *In vivo* kinetics of primary tumor growth in MDA-MB-231 and MMTV-WNT1 models. **(F)** Matched Odyssey and immunohistochemistry (IHC) of representative orthotopic BC (first row), intra venous BC (second row), intra-splenic CRC (third row), and spontaneous MMTV-WNT1 BC (fourth row) liver slices. ICG signal is represented as black spots in the white background for Odyssey images. Anti-human nuclei signal (brown nuclei) is represented in the IHC images of orthotopic BC, intra-venous BC, and intra-splenic CRC models. Hematoxylin-eosin signal is represented in the IHC image for MMTV-WNT1 model. **(G)** Hematoxylin-eosin representative image of MMTV-WNT1 primary tumor. **(H)** Matched Odyssey and IHC images of liver slices where no ICG signal was detected.

**Fig. 2:**
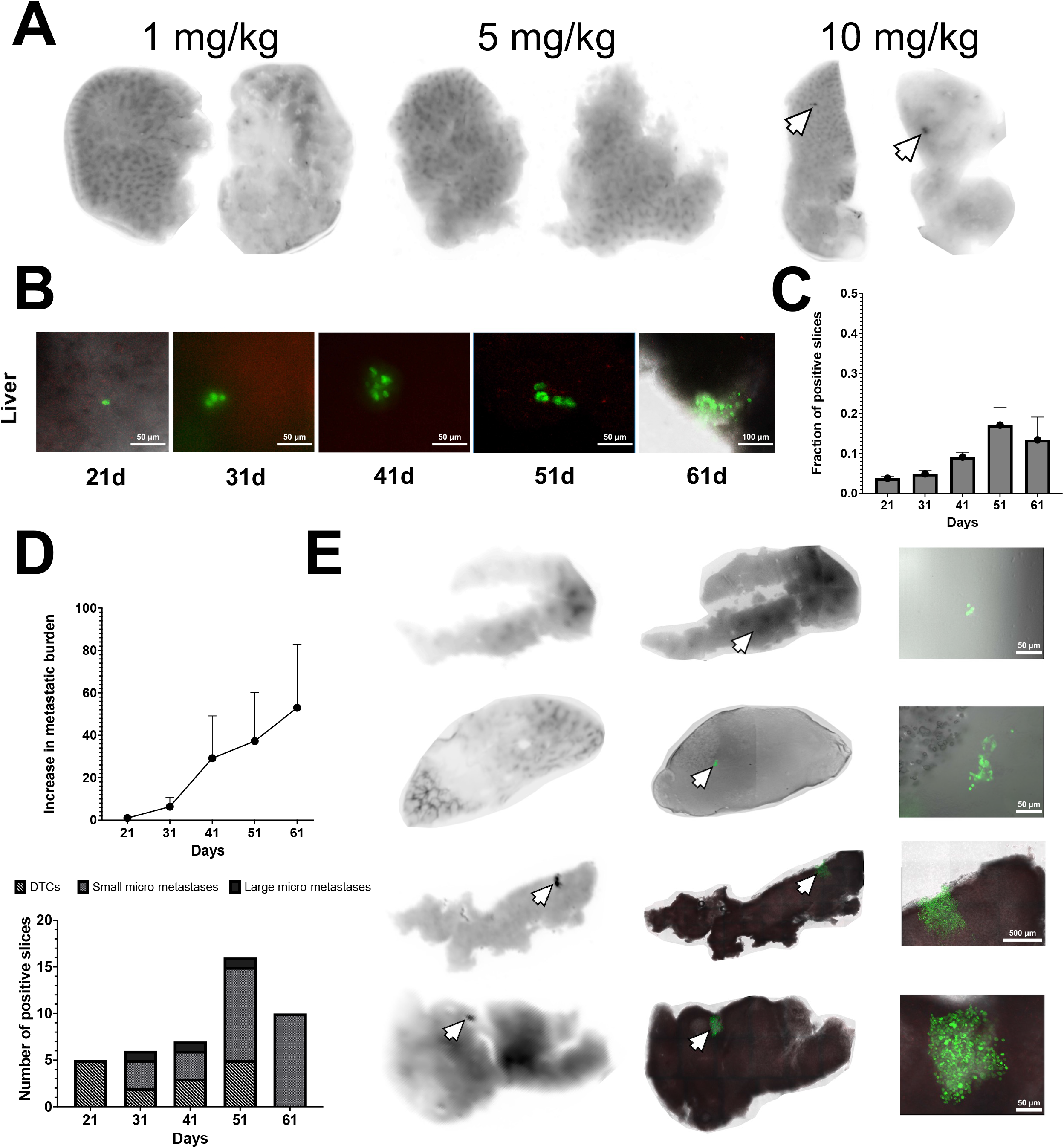
Optimization of ICG-sensitivity. **(A)** Representative Odyssey images of liver slices (intravenous BC model) where ICG signal was investigated flowing administration of different ICG dosage. **(B)** Representative fluorescence microscopy images of liver disseminated tumor cells and micro-metastases over time (orthotopic BC model). **(C)** Fraction of liver slices in which either disseminated tumor cells or micro-metastases were detected at indicated time points. **(D)** Kinetics of increase in liver metastatic burden (orthotopic BC model) normalized to the first time point (i.e., 21d after BC transplantation). Metastatic burden was measured as the fraction of the area occupied by metastatic cells over the total area of all liver slices. **(E)** Matched Odyssey images and fluorescence microscopy images of liver slices (61d post orthotopic injection) displaying single disseminated tumor cells, clusters of disseminated tumor cells, and micro-metastases (orthotopic BC model).

In the intra-splenic transplantation, 10^6^ U11 CRC PDX cells were resuspended in 200 μl of PBS (Thermo Fisher Scientific) and injected in the exposed spleen of 8–12-week-old NSG mice. Mice were sacrificed 10 days after injection to inspect liver micro-metastases.

In the spontaneous breast cancer MMTV-WNT-1 model, only mice displaying clearly palpable tumors at 5-6 months of age were selected for further investigation. Tumor size was weekly measured by caliper and masses were allowed to grow up to 1-1.5 cm^3^. Then, mice were sacrificed to inspect liver metastases.

Independently of the setting, mice received an intra-peritoneal injection of ICG (10mg/kg) 16 hours before sacrifice.

### LI-COR Odyssey IR tissue scan

In all the *in vivo* settings investigated, liver was carefully harvested so to maintain its integrity and the gallbladder removed. Collected liver was placed in cold PBS and maintained on ice till sectioning. An identical protocol was adopted to harvest mouse lungs.

To generate organ slices with thickness ranging from 0.5 to 1 mm, the liver was divided into the five constituting lobes and each lobe glued on the stage of a Leica VT1200 vibratome. When the glue dried out, the stage was covered with cold PBS and the lobe cut into slices. Instead, to generate slices with thickness ranging from 0.15 to 0.25 mm, liver was divided into the five constituting lobes and each lobe included in 4% low melting agarose (55°C) in PBS. Lobe inclusion was performed on ice until solidification. Then, the whole block of agarose was glued on the stage of a Leica VT1200 vibratome. When the glue dried out, the stage was covered with cold PBS and the lobe cut into slices. To generate lung slices with thickness ranging from 0.1 to 0.25 mm, the same protocol was adopted.

Slices of interest were imaged with LI-COR Odyssey CL-x platform. To this aim, slices were places into the machine and the 800 nm wavelength selected to image uniquely near-infrared signal. We performed high quality imaging in the DNA-gel Preset, setting the image resolution at 21 µm and the intensity at its lowest value (L2.0). Upon imaging, slices of interest underwent further processing.

### Immunohistochemistry and fluorescence microscopy

To assess whether ICG spots correspond to bona fide human cells, slices containing ICG spots were fixed for 2h in 4% formaldehyde (Sigma-Aldrich) and subsequently transferred to 70% ethanol overnight (Panreac Applichem) prior to paraffin inclusion. Paraffin-embedded slices were cut into 5μm thick sections which were subsequently stained with both hematoxylin-eosin and antihuman nuclei (Catalog # RBM5-346-P1) antibody (2μg/ml). Stained sections were subsequently imaged with a Leica DM6 B HistoFluo microscope.

To assess whether ICG signals overlapped with H2B-GFP enriched lesions, lung and liver slices with thickness ranging from 0.15 to 0.25 mm were imaged with a Nikon Eclipse TI2-E (inverted) microscope. Slices were screened to identify uniquely those which displayed metastatic colonization (single disseminated tumor cells, small micro-metastases (<10 cells), large micro-metastases (>10 cells), and macro-metastases (>500 cells were considered for this analysis). Selected slices were imaged with the LI-COR Odyssey as described above.

### Manual coring of ICG spots

To core ICG positive spots, we took advantage of glass capillaries with 1mm diameter. Liver slices displaying an ICG spot were set apart and imaged for a second time to clearly determine the spot position in the slice. Then, we manually cored the region which contained the spot and dispensed the cored region in a cold PBS-containing 1.5 ml Eppendorf tube. The cored region was resuspended in cold PBS and cells disaggregated by pipetting. In the meantime, another image of the slice was taken so to ensure that the ICG spot was completely removed. Viability of cored cells was checked in an independent experiment after 30 minutes on ice via FACS: the percentage of dead cells was computed by staining samples with propidium iodide (250 μg/ml). Cored regions were then processed for scRNA-seq within 10 minutes from coring.

### scRNA-seq analysis

Cored liver regions and matched primary tumors (PTs) underwent scRNA-seq. Tissue fragments were processed as described above, while matched PTs were digested following the protocol from the Miltenyi human tumor dissociation kit (Prior to sequencing, cells were resuspended in cold PBS and counted in a Burker chamber. Then, cells were pelleted, the supernatant removed, and the pellet resuspended in PBS = 0.5% FBS at the final concentration of 500-1000 cells/μl. Single-cell suspension was mixed with reverse transcription mix using the Chromium Single-cell 3’ reagent kit protocol V3.1 (10x Genomics) and loaded onto 10x Genomics Single Cell 3’Chips to generate single-cell beads in emulsion. The gel beads are coated with unique primers bearing 10× cell barcodes, unique molecular identifiers (UMI) and poly(dT) sequences. The chip was then loaded onto a Chromium instrument where RNA transcripts from single cells are reverse-transcribed. cDNA molecules from one sample were pooled, amplified, and the amplified cDNAs were fragmented. Final libraries were generated by incorporating adapters and sample indices compatible with Illumina sequencing and quantified Qubit (ThermoFisher Scientific). The size profiles of the sequencing libraries were examined by Agilent Bioanalyzer 2100 using a High Sensitivity DNA chip (Agilent). Two indexed libraries were equimolarly pooled and sequenced on Illumina NOVAseq 6000 platform using the v1.5 Kit (Illumina) with a customized paired end, dual indexing (26/8/0/91-bp) format according to the manufacturer’s protocol. A coverage of ∼50.000 reads per cell was adopted for each sequencing run.

The raw Illumina BCL files obtained from sequencing of the three samples were demultiplexed and converted to FASTQ format using the Illumina *bcl2fastq* software (v2.20.0.422). *FastQC* (v0.11.5) https://www.bioinformatics.babraham.ac.uk/projects/fastqc/ was used to evaluate the quality of sequenced reads. Reads were then aligned to the GRCh38 human reference genome using 10X Genomics *Cell Ranger count* (v6.1.2), exploiting the GENCODE gene set GTF file (v32, version: 2020-A) for the annotation of the genes. The program, which only includes reads which are confidently mapped, non-PCR duplicates, containing valid barcodes and UMIs, finally quantifies the expression of transcripts in each cell and generate the gene expression matrix of number of UMI for every cell and gene, which is the final output of the pre-processing part. All the further steps of the analysis were performed in R (v4.2.2) https://www.r-project.org/ using the Seurat package (v4.2.1), unless otherwise specified. Each 10x library, henceforth referred to as ‘sample’, was individually quality checked, and cells were filtered to ensure good gene coverage, a consistent range of read counts and low numbers of mitochondrial reads, applying common quality metrics. Specifically, cell and gene quality control (QC) were performed for each sample separately. For each sample, and cells with QC metrics (i.e., number of UMIs and number of genes detected per cell) falling outside of the [median – 5*M.A.D., median + 5*M.A.D.] range were discarded. Additionally, cells with cell complexity > 0.8 (the ratio between the log10 of number of genes per cell and the log_10_ of the number of UMIs per cell) and mitochondrial genes percentage > 20% was filtered out. After filtering steps, samples were merged using merge() function while sctransform (using the glmGamPoi method, from the glmGamPoi package (v1.10.0)) with default parameters is used to normalize, scale the counts and to identify the most variable genes (n=3,000) of the merged Seurat object. Moreover, cellCycleScoring() function in Seurat is used to assign each cell to a cell cycle phase while in order to identify population that are present across datasets, samples has been integrated following the workflow suggested by Seurat and UMAP graphical representation has been produced employing in sequence the functions runPCA and runUMAP, in order to visualize cells in a two dimensions plot. After dimensionally reductions steps, the clustering approach has been conducted using *FindNeighbors()*and *FindClusters()*employing several resolution parameters (0.1, 0.3, 0.5, 0.7 and 0.9) to then proceed with resolution= 0.5 that allow to identify 12 clusters. Finally, FindMarker () function from Seurat has been used to determine difference among Seurat clusters or samples (metastases vs PT). Marker genes were used to identify over-represented pathways in the clusters or samples of interest. To this aim, the top 50 marker genes (when available) were used to interrogate the Gene Ontology (GO) and Hallmark collections of gene sets in the Molecular Signature Database (MSigDB). The top 50 over-represented collection per cluster (or sample) were considered and used to establish functional gene signatures enriched in each cluster (or sample) of interest.

### Statistical analysis

All the ranges reported in the text correspond to mean ± standard deviation. Bar plots and line plots display mean ± standard deviation, functional enrichment plots display the false discovery rate (FDR)value associated with the plotted pathway or gene. Volcano plots were generated through GraphPad software by plotting the log_2_FC vs -log_10_(FDR) of the genes that were identified as differentially expressed (considering the p-value) in the micro-metastases vs PTs. The Kaplan-Meier curve depicted in Fig. 4H was generated through the online available tool Kaplan-Meier plotter (https://kmplot.com/analysis/index.php?p=service). Five-year overall survival was considered as clinical outcome of interest. Log-rank test p-value is reported in the corresponding figure.

## RESULTS

### ICG staining allows visualization of liver micro-metastases

To assess whether ICG allows for the labeling of liver micro-metastases, we investigated early liver colonization using different cancer mouse-models, including the human MDA-MB-231 triple negative BC cell line, the U11 human colorectal cancer (CRC) patient-derived xenograft (PDX), and the BC MMTV-WNT-1 transgenics (**21**).

MDA-MB-231 cells were transplanted in NSG mice either orthotopically (transplantation in the ninth mammary gland; Fig. 1A) or systemically (direct intra-venous injection; Fig. 1B). 10^6^ U11 CRC cells were transplanted in NSG mice via intra-splenic injection, to mimic the most frequent route of CRC metastatization (Fig. 1C). Ultimately, MMTV-WNT-1 mice, adopted to explore cancer-cell seeding in the context of spontaneous tumorigenesis, displayed a ∼30% disease-penetrance at 5-6 months of age (Fig. 1D), in line with previous studies (**21**).

Liver colonization was assessed at: i) ∼40 days upon orthotopic implantation of MDA-MB-231 cells, when tumor masses reached the maximum permitted size (Fig. 1E); ii) at 28 days after intravenous injection of MDA-MB-231 cells; iii) at 10 days after intra-splenic injection of U11 CRC cells; and iv) at 5-6 months of age of the MMTV-WNT-1 mice, when tumor size reached the maximum permitted size (Fig. 1E). Prior to sacrifice, mice were intra-venously injected with ICG (10mg/kg), 16 hours before organ collection. Upon mouse sacrifice, livers were harvested, fresh tissue cut with vibratome (slice thickness ranging from 0.5 to 1 mm) and slices processed with the LI-COR Odyssey platform. We observed intense ICG spots in the liver parenchyma of all cancer-cell injected mice, regardless of the model (Fig. 1F).

To investigate correspondence in the liver between ICG signal and *bona fide* micro-metastases, spot-containing liver slices were fixed, paraffin-included and stained with anti-human nuclei antibodies or hematoxylin eosin to detect, respectively, MDA-MB-231 and U11 PDX cells or MMTV-WNT-1 BC cells. We scored consistent overlaps between ICG spots and anti-human signals in all human models (Fig. 1F), or areas with ductal structures resembling those of the corresponding primary tumors in the MMTV-WNT-1 model (Fig. 1G). Of note, immunohistochemistry (IHC) staining of liver slices devoid of spots revealed no presence of tumoral cells (Fig. 1H).

Thus, ICG can be used to successfully mark liver micro-metastases across multiple *in vivo* tumor models and experimental conditions.

### Optimization of ICG-sensitivity

To optimize our ICG assay, we next investigated the effects of ICG dosage and liver-slicing thickness on the sensitivity of micro-metastases identification. In previous clinical studies, ICG was intra-venously administered at 0.5 mg/kg (**22,23**) and signals investigated in patients at different time-points ranging from ∼24 to >48 hours. In pre-clinical settings, ICG was retro-orbitally administered at 2.5 - 5 mg/kg (**18,24**) and signal investigated 12-24 hours after. We administered three different ICG concentrations (1, 5, and 10 mg/kg) 28 days after intra-venous injection of MDA-MB-231 cells and imaged the ICG signal 16h after. Upon sacrifice, livers underwent vibratome cutting (slice thickness ranging 0.1 - 1 mm) and imaging with the LI-COR Odyssey imaging platform. As shown in Fig 2A, only the highest ICG dose (10 mg/kg) yielded detectable ICG spotting.

To investigate the relevance of slice thickness, we used MDA-MB-231 engineered with the green fluorescent protein (GFP) linked to the histone 2B protein (H2B-GFP reporter; **25**). Upon H2B-GFP lentiviral infection and fluorescence-based sorting, GFP-expressing MDA-MB-231 cells were orthotopically transplanted in the ninth mammary gland of NSG mice. Hepatic colonization was monitored every ten days (n=2 mice per time point), starting 21 days after transplantation and ending 61 days after transplantation. Mice were then intravenously injected with 10 mg/kg ICG and sacrificed after 16h. Livers were collected and cut in slices with thickness ranging 0.15-0.25 mm. Given the high number of slices (∼50 per organ), slices were first screened by immunofluorescence to identify positive slices (i.e., slices harboring H2B-GFP positive cells). Interestingly, liver was already colonized at 21 days after transplantation by disseminated tumor cells (DTCs; Fig. 2B) which appears as isolated cells or small groups of few cells. Over time, we detected an increase in liver colonization, resulting in a higher fraction of positive slices (Fig. 2C) and in larger ICG-positive groups of cells with a higher fraction of liver surface occupied by metastatic cells (Fig. 2D). While DTCs were still present at later time points (Fig. 2E), small (<50 cells) and large micro-metastases started to populate the liver parenchyma, suggesting a kinetic of colonization for which only a fraction of DTCs proliferate enough to progress to larger lesions and/or PTs constantly produce DTCs regardless of age or size.

The micro-metastasis-containing slices were then analyzed with the LI-COR Odyssey platform. We found that ICG spots were present uniquely in correspondence of large micro-metastases (Fig. 2E), while DTCs or small micro-metastases were not detected (Fig. 2E).

Thus, optimal detection of liver micro-metastasis by ICG staining requires a minimum dosage of intravenously injected ICG (10 mg/kg) and a minimum micro-metastasis cellularity (>50 cells), while tissue slicing can be scaled down from 1 mm to hundreds of microns.

### ICG staining allows visualization of lung micro-metastases and monitoring of treatment responses

We then asked whether our ICG assay can be used to identify other metastatic sites, such as lung, and to quantify the effect of anti-metastatic treatments. We initially investigated by fluorescence the kinetics of lung colonization by MDA-MB-231 cells after orthotopic transplantation (same time-points as in Fig. 2B-D). As shown in Fig. 3A, H2B-GFP MDA-MB-231 cells infiltrate lungs as single cells at early time-points (i.e., 21d), to then progress into micro- and, eventually, macro-metastases (Fig. 3B). To test for their anti-metastatic potential, we screened *in vitro* several inhibitors of cellular oxidases that affect cellular redox status, including allopurinol, apocynin, diphenylene iodonium, azalastan and others. We selected safinamide, an inhibitor of monoamine oxidase B that is primarily used to treat Parkinson’s disease (**20**) and more recently shown to possess potential anti-metastatic activities (**26**). Safinamide did not significantly alter proliferation of MDA-MB-231 cells *in vitro* (Fig. 3C). To track *in vivo* metastatization and response to treatment, we generated two distinct subpopulations of MDA-MB-231 (H2B-GFP and H2B-RFP), which were treated (H2B-RFP cells) or not (H2B-GFP cells) for 72 hours *in vitro* with 10 μg/ml safinamide. The two subpopulations were mixed 1:1 and intra-venously injected in NSG mice. Lungs were collected 28 days post-injection and 16 hours after intra-venously administration of 10 mg/kg ICG, cut into 0.5 mm slices, and scanned with the LI-COR Odyssey, which revealed the presence of ICG spots in 100% lung slices from each animal (representative spots are shown in Fig. 3D). Imaging of the same slices by fluorescence microscopy showed a strong correspondence between ICG accumulation, nuclear fluorescence, and anti-human antibody signal (Fig. 3E). As observed in the liver, only large size micro-metastases (>50 cells) were successfully identified by ICG (Fig. 5D-E). Importantly, the ICG spots colocalized with anti-human nuclear staining, as revealed by immunohistochemistry (Fig. 3F). Strikingly, no H2B-RFP signal was scored by immunofluorescence in any of the positive slices, where the ICG signal overlapped with H2B-GFP positive cells (Fig. 5G). Thus, ICG staining can be efficiently exploited to mark lung micro-metastases and monitor treatment responses or highlight selected foci in competitive transplantation assays.

**Fig. 3:**
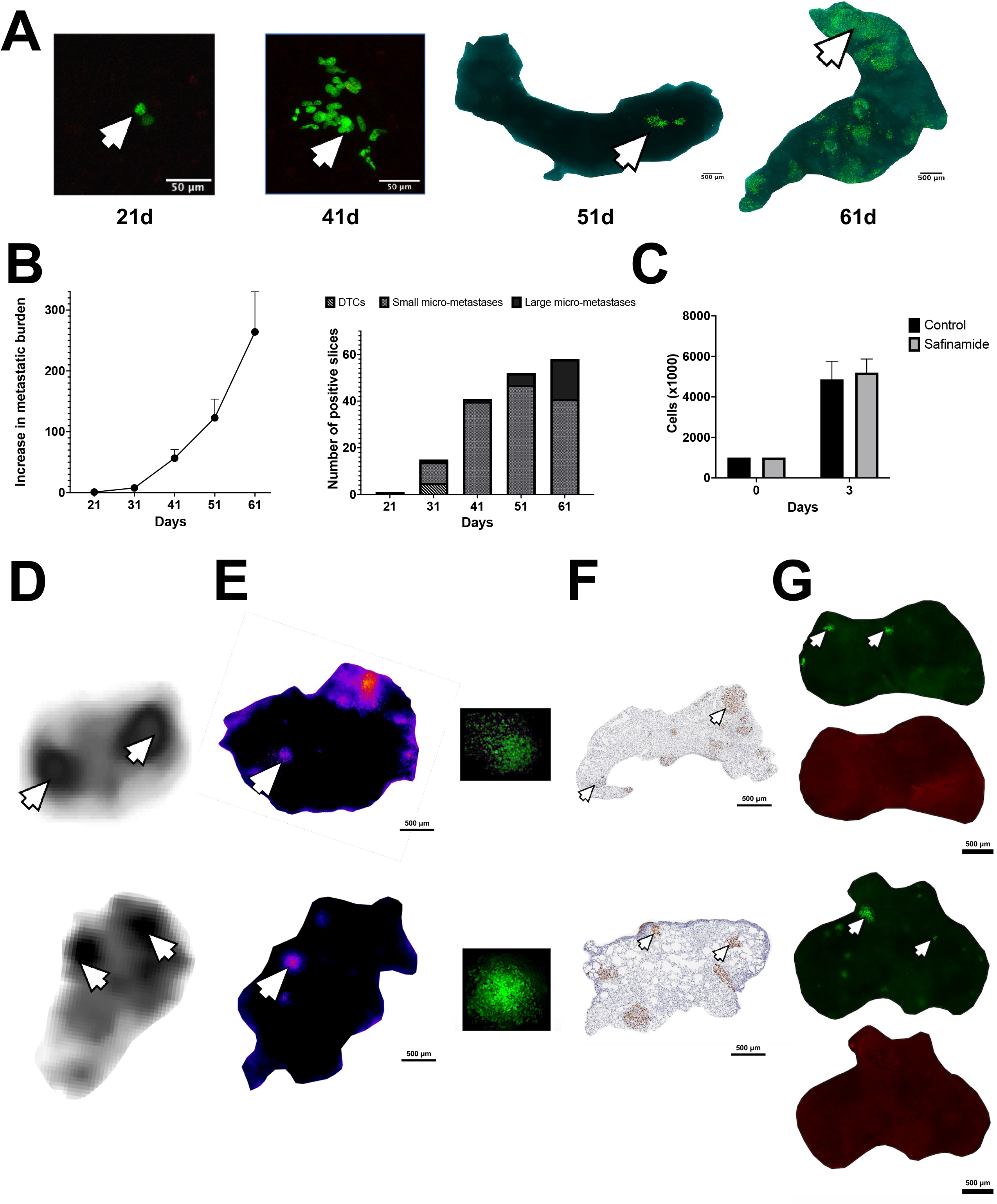
ICG staining allows for the visualization of lung micro-metastases and monitoring of treatment responses. **(A)** Representative fluorescence microscopy images of lung disseminated tumor cells, micro-, and macro-metastases over time (orthotopic BC model). **(B)** Kinetics of increase in lung metastatic burden (orthotopic BC model) normalized to the first time point (i.e., 21d after BC transplantation). Metastatic burden was measured as the fraction of the area occupied by metastatic cells over the total area of all lung slices. **(C)** MDA-MB-231 cell count immediately after plating and after 72h of safinamide treatment at 10μg/ml. **(D)** Representative Odyssey images of lung slices (orthotopic BC model). ICG signal is represented as black spots on the white background. **(E)** Matched fluorescence microscopy and Odyssey images of lung slices displaying micro-metastases (orthotopic BC model). **(F)** IHC images of lung slices matched to images shown in (E). Anti-human nuclei signal (brown nuclei) was investigated. **(G)** Representative fluorescence microscopy images of lung slices infiltrated by either untreated (H2B-GFP) or *in vitro* treated (10μg/ml safinamide; H2B-mCherry) MDA-MB-231. The same slices were investigated to determine the presence of both cell types.

**Fig. 4:**
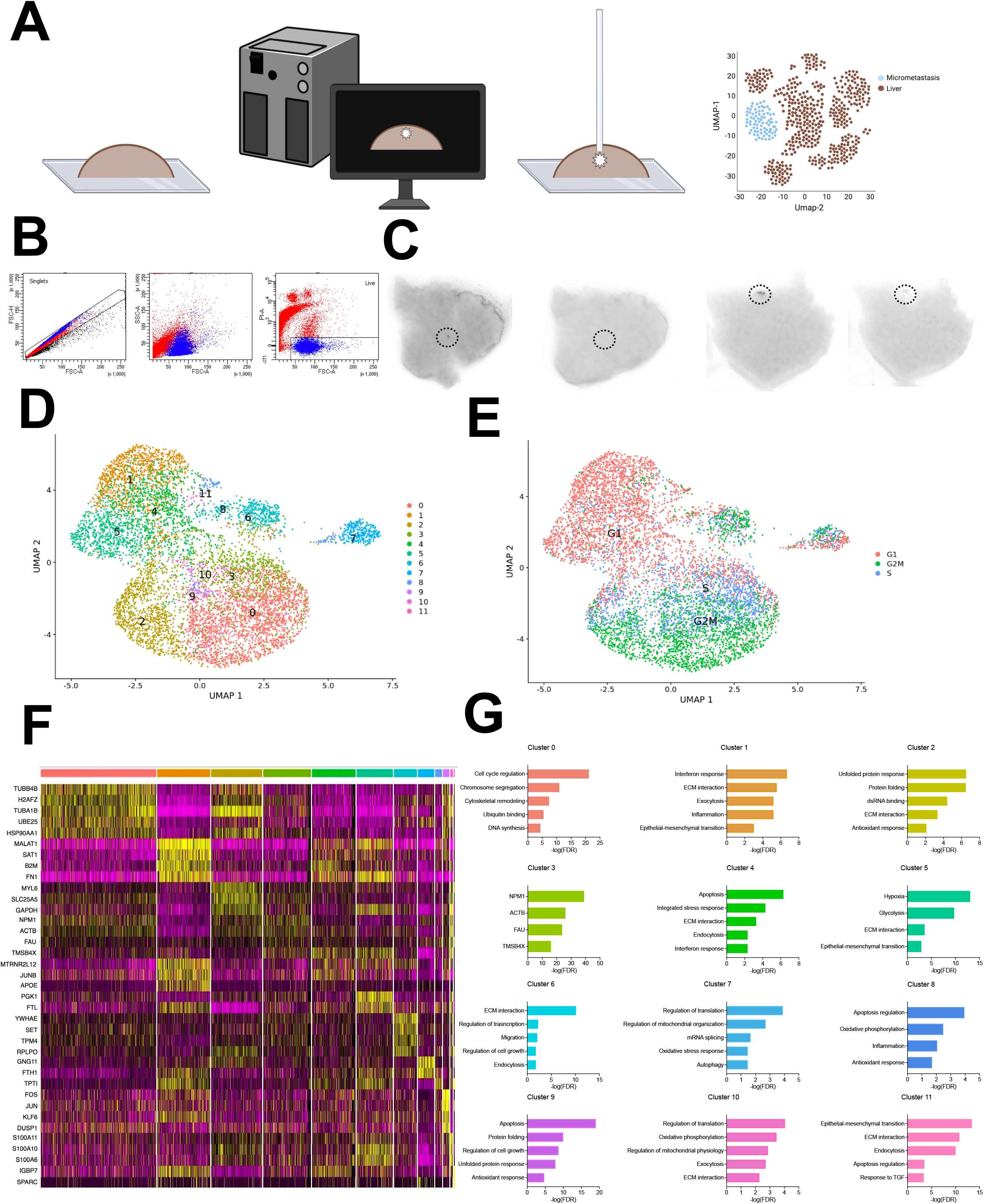
Recovery of micro-metastasis for single-cell transcriptional analyses. **(A)** Schematic representation of the coring protocol for the single-cell profiling of orthotopic BC model micro-metastases. **(B)** FACS analysis of representative liver single-cell suspensions 30 minutes after coring. Cells were gated to discriminate singlets (FSC-A vs FSC-H) and alive cells (FSC-A vs PI-A). **(C)** Representative Odyssey images of liver slices (orthotopic BC model) before and after manual coring. **(D)** UMAP of 7,594 BC cells (7,456 from primary tumors and 138 from matched micro-metastases; orthotopic BC model) colored by cluster. **(E)** UMAP of 7,594 BC cells (7,456 from primary tumors and 138 from matched micro-metastases; orthotopic BC model) colored by cell cycle score. **(F)** Marker gene analysis of the 11 clusters. The top 5 marker genes per cluster are shown, when available. Color code represents the level of expression of each gene (from yellow-high to violet-low). **(G)** Bar plot representing the functional enrichment terms of the enriched gene sets across the 11 clusters. For each term, the FDR is plotted (for cluster 3 we show FDR of the 4 identified marker genes).

**Fig. 5:**
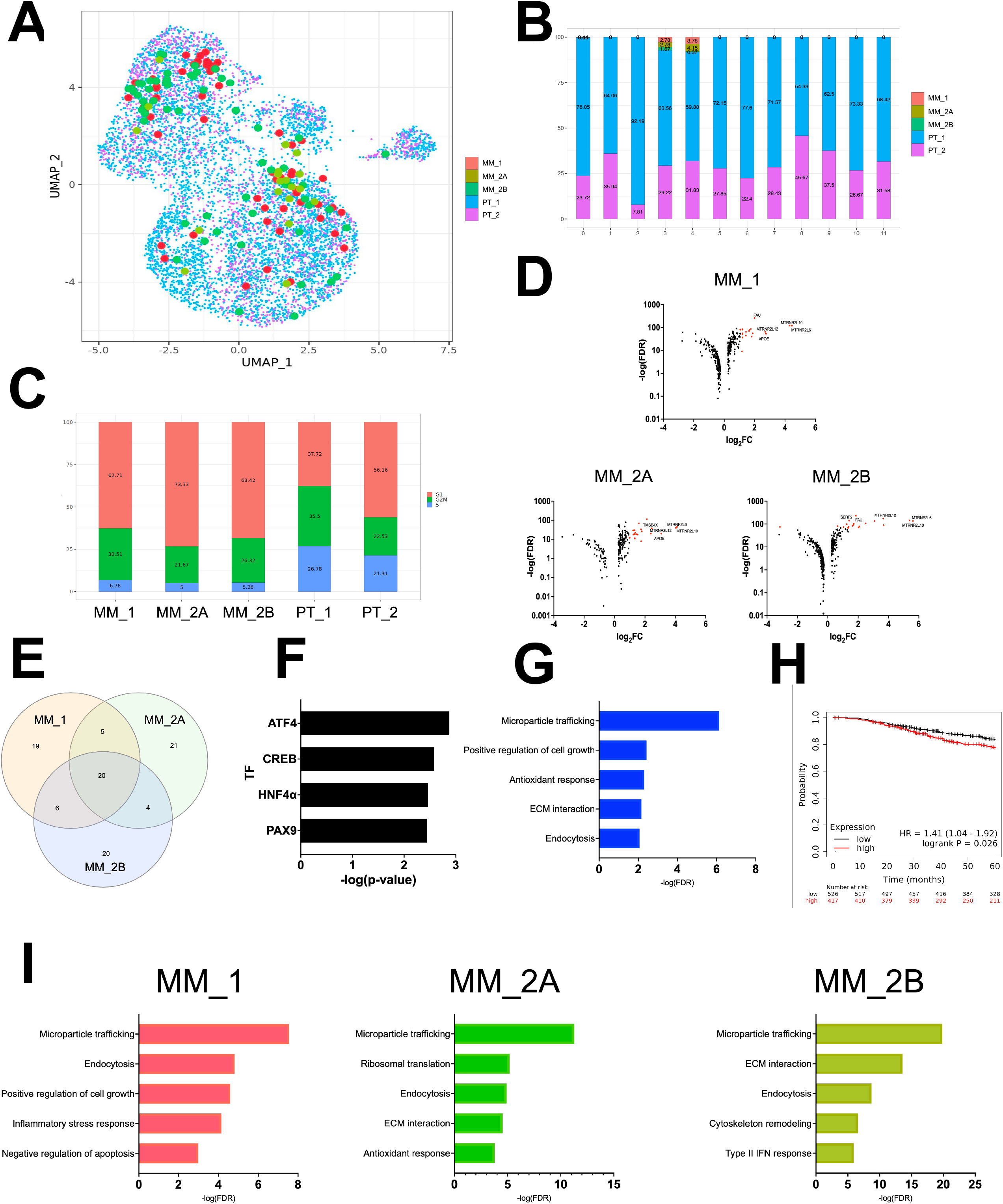
ICG-identified lesions upregulate shared and private pathways. **(A)** UMAP of 7,594 BC cells (7,456 from primary tumors and 138 from matched micro-metastases; orthotopic BC model) colored by sample. Micro-metastasis cells are represented with larger dots. **(B)** Stacked bar plot of cluster composition in terms of samples. **(C)** Stacked bar plot of cell cycle distribution of each sample. **(D)** Volcano plot representing DEGs in the three micro-metastases. For each DEG, we reported the corresponding logFC (x-axis) and the -log(FDR) (y-axis). The top 20 upregulated DEGs (ranked by FDR) are colored in red. **(E)** Venn diagram representing the top 50 upregulated DEGs in each micro-metastasis. **(F)** Bar plot of the transcription factors (TFs) identified by the PASTAA algorithm considering the 20 common DEGs upregulated across the three micro-metastases. **(G)** Bar plot representing the functional enrichment terms of the enriched gene sets identified by using the 20 common DEGs upregulated across the three micro-metastases. For each term, FDR is plotted. **(H)** Survival analysis of the 5-year of patients according to the expression of the 20 common DEGs upregulated across the three micro-metastases. **(I)** Bar plot representing the functional enrichment terms of the enriched gene sets identified within each micro-metastasis. For each term, FDR is plotted.

### Recovery of micro-metastasis for single-cell transcriptional analyses

We then explored the possibility of collecting ICG spots for scRNA-seq analyses. Mouse livers were harvested after orthotopic injection of MDA-MB-231 BC cells (same time points as in Fig. 1A) and cut it into 0.5 mm slices (Fig 4A), a thickness allowing scanning of the whole organ in less than <10 minutes, thus preserving its integrity, which is rapidly lost in 30 minutes on ice (Fig. 4B). Slices with clear ICG spots were identified by the LI-COR Odyssey imaging and manually cored using 1mm-size glass capillaries (Fig. 4C). Cells from the tissue core were resuspended in PBS supplemented with 10% FBS, counted, and to a concentration of 500-1000 cells/ µl for scRNA-seq analyses.

We obtained high-quality scRNAseq data from 3 micro-metastases (59, 19, and 60 cells in MM_1, MM_2A, and MM_2B, respectively) and two matched PTs (5,485 and 1,971 cells in PT_1 and PT_2, respectively). While tumor cells were predominant in the PT samples (>95%), not surprisingly they were extremely rare in the micro-metastases, ranging from 0.87% to 1.81%.

The 7,594 tumor cells of our dataset (7,456 from PTs and 138 from matched micro-metastases) were grouped in 11 stable clusters (Fig. 4D). Analyses of cell-cycle distribution by the Seurat package showed enrichment in cycling cells in Clusters 0, 2, 6, and 9 (% S-G2M cells: 82.04 ± 11.38), whereas the remaining clusters were largely enriched in non-dividing cells (% of G1 cells: 69.66 ± 20.61) (Fig. 4E). The top 50 marker genes *per* cluster were used to identify over-represented gene-sets in the MSigDB (Gene Ontology (GO) and Hallmark collections) (Fig. 4F, Suppl. Table 1). Cluster 3 presented with only 4 marker genes, thus preventing gene-set enrichment analyses. Of the highly proliferating clusters, only Cluster 0 was enriched for terms related to cell growth (Fig. 4G). Clusters 6 and 10, instead, were mainly enriched for terms related to gene expression regulation, interaction with the extracellular matrix (ECM), and exo/endocytosis. Intriguingly, ECM interaction was strongly upregulated also in Cluster 11, which is mainly enriched for terms related to the migratory phenotype. Ultimately, remaining clusters showed upregulation of different stress response pathways, including unfolded protein response (UPR; Clusters 2 and 9), Interferon response (Clusters 1 and 4), Hypoxia response (Cluster 5), Oxidative stress response (Clusters 7 and 8) and Response to inflammation (Clusters 1 and 8). Of note, stress response pathways were associated to the regulation of apoptosis (Cluster 4, 8, and 9) and autophagy (Cluster 7). Clusters 8 and 10 showed upregulation of oxidative phosphorylation, while Cluster 5 was mainly glycolytic. Interestingly, our dataset displays a transcriptional heterogeneity which largely resembles what was reported in previous studies in BC (**27**). Therefore, ICG can be successfully exploited to guide manual coring of vital hepatic micro-metastases which can be directly analyzed by scRNA-seq.

### ICG-identified lesions upregulate shared and private pathways

We ultimately focused on micro-metastases, to identify distinguishing genes and pathways (Suppl. Table 2). We found that micro-metastases populate the less proliferative Clusters (accounting for 7.23% and 8.3% of cells in Clusters 3 and 4 respectively; Fig. 5A and 5B) and, consistently, are mainly composed of non-cycling cells (68,15 ± 5,31% of cells in G1 phase; Fig. 5C). We performed differential gene-expression analyses by comparing each micro-metastasis to the pool of the two PTs. This analysis revealed an average of 150 significantly (FDR < 0.05) upregulated genes per each micro-metastasis (150 ± 28) (Fig. 5D), with a large degree of overlap across samples: 20 shared among three and 15 among two micro-metastasis. ∼20 differentially-expressed genes (DEGs) were instead specific to each micro-metastasis (Fig. 5E).

We initially focused on the 20 common genes searching for putative upstream transcription-factors (TFs), leveraging the online tool PASTAA (**28**), and identified 4 TFs (Fig. 5F) associated to the regulation of the integrated stress responses (ATF4, **29**), cell proliferation (CREB, **30**), antioxidant response (HNF4, **31**), and ECM interaction (PAX9, **32**). Consistently, GSEA analyses using the 20 common genes showed upregulation of terms related to microparticle trafficking, endocytosis, positive regulation of cell growth, interaction with the ECM, and antioxidant response (Fig. 5G). Remarkably, higher expression of the 20 common genes in BC patient primary tumors predicted worse 5-year overall survival (p=0.026; Fig. 5H).

We then performed GSEA analyses using the top 50 DEGs of each micro-metastasis separately. We found that each micro-metastasis upregulates terms related to microparticle trafficking and endocytosis (Fig. 5I). However, each micro-metastasis displayed a unique subset of upregulated pathways: stress response and positive regulation of cell growth in MM_1; ribosomal translation and antioxidant response in MM_2A; interferon response and cytoskeleton remodeling in MM_2B (Fig. 5I). Notably, ECM interaction was present in both MM_2A and MM_2B.

Therefore, our results revealed that BC micro-metastases upregulate a common subset of genes and pathways, while at the same time being enriched for specific transcriptional traits that are not shared with the other lesions.

## DISCUSSION

Metastatic progression represents the deadliest outcome of most tumors, including BC. Unfortunately, a relevant fraction of patients displays clinically undetectable metastases at the time of diagnosis, which therefore jeopardize the administration of drugs aimed to reduce metastatic spreading. In this scenario, a deep comprehension of the molecular phenotype of clinically undetectable micro-metastases becomes of primary importance to tailor therapeutic strategies aimed at blocking micro-to macro-metastatic progression.

Data from sizeable, millimeter range, metastatic lesions in preclinical models or in patients suggest that critical steps for disease manifestation occur after the engraftment of disseminating cells within the target organ (**33**). These steps require interactions with the microenvironment and include long term quiescence and phenotypic remodeling (**34,35**) that are poorly detailed in small aggregates of infiltrated tumor cells. Furthermore, single cell transcriptomics of small metastatic lesions has been obtained by FACS sorting, which necessarily led to the loss of lesion structural organization (**36**) and then to the loss of all the phenotypic information regarding molecular pathways simultaneously activated in each lesion. Therefore, a method coupling the isolation of small micro-metastases to the transcriptional investigation of the whole lesion at single cell level is extremely relevant and needed.

ICG represents a widely used contrast agent in surgical oncology (**37**), however the molecular mechanisms of ICG binding to tumor lesions are still under debate. While several studies pointed out that tumor cells are stained because of an increased uptake and retention when compared with normal cells (**38**), other studies suggested that ICG binds extracellular matrix components like collagen and fibronectin (**39**). In this latter case, the denser tumor ECM accounts for the increased ICG signal in cancer cells when compared to the surrounding parenchyma. Our results cannot rule out any of the two hypotheses. Indeed, we showed that ICG binds *bona fide* tumor cells in the liver parenchyma, generating signal spots that require a minimum cellularity to be detected. However, this requirement may result from the necessity to accumulate enough ICG either within cells or in the extracellular matrix.

Our findings revealed that BC metastatic cells infiltrate the liver parenchyma as single cells and slowly proliferate to small micro-metastases in the niche. These micro-metastases rarely reach a sufficient size to be identified through ICG screening, while most of them remain dormant or progress till small cellular aggregates (<100 cells indicatively). This result suggests that most liver micro-metastases do not progress from single disseminated tumor cells to micro- and macro-metastases, but rather they undergo dormancy and persist in the hepatic parenchyma as single cells or small groups.

We managed to manually core BC liver micro-metastases and to successfully analyze them by scRNA-seq, demonstrating that ICG can be successfully exploited to guide the coring of small micro-metastases for subsequent single-cell transcriptional profiling. Our results revealed that micro-metastases over-express a specific subset of genes when compared with matched PTs. These genes are mainly involved in microparticle trafficking, endocytosis, positive regulation of cell growth, interaction with the ECM, and antioxidant response. This finding suggests an active transcriptional phenotype of micro-metastatic lesions, by which tumor cells may adapt to the surrounding microenvironment and accomplish the transition towards macro-metastases. Indeed, genes involved in the antioxidant response and microparticle trafficking can be activated to prevent Kupffer cell-mediated oxidative stress (**40,34**) and to polarize liver macrophages towards a cancer-permissive environment (**41**). On the other hand, the activation of the endocytic pathway was previously reported to be crucial for the metastatic progression of several cancer types (**42**). Indeed, this process allows for continuous remodeling of the interface between tumor cells and ECM and favors the invasion of the surrounding tissue. Therefore, our data suggest that BC micro-metastases can condition the liver parenchyma by inducing a cancer-permissive microenvironment where they can proliferate and migrate. Importantly, genes involved in these pathways were found to statistically predict worse BC patient survival, therefore corroborating the clinical relevance of our findings. Importantly, albeit investigated micro-metastases displayed a common core of upregulated genes, our approach allowed to define the inter-metastases transcriptional heterogeneity, which could never be investigated through approaches that destroy the structure of the micro-metastasis itself. We found that each micro-metastasis harbors a subset of private upregulated genes, which can be involved in either inflammatory stress response, ribosomal translation, or interferon response. Of note, we previously demonstrated that the hyper-activation of interferon-related genes is fundamental for the successful accomplishment of the metastatic cascade (**27**). Our results suggest that the activation of these genes support the transition from micro-to macro-metastasis.

Eventually, we revealed that ICG can be successfully adopted to identify BC micro-metastases in the lung parenchyma. Lungs represent a privileged metastasis site in triple-negative BC (**3**), thus advocating for drugs that efficiently inhibit lung colonization. MAO inhibitors have recently shown efficacy in dampening the metastatic progression of prostate cancer (**43**), thereby laying the groundwork for a possible exploitation in other tumor settings. By coupling ICG staining and fluorescence microscopy, our data indicate that *in vitro* treatment of metastatic BC cells with the MAO-B inhibitor safinamide leads to a complete abrogation of the metastatic phenotype. Further studies are however necessary to determine whether safinamide administration can represent a feasible therapeutic strategy in BC.

Therefore, our results revealed an unprecedented diagnostic power for ICG, which can be successfully exploited to identify small size micro-metastases and perform both molecular and functional studies.

## ACKNOWLEDGEMENTS

We thank Mario Varasi for helpful discussion and Stefania Averaimo for the editing of the manuscript.

## AUTHOR CONTRIBUTIONS

NR and VG performed cellular and *in vivo* experiments. MS performed the *in vivo* experiments and the tissue analysis. GP and FC performed the computational analysis. SR performed imaging experiments. NR, VG, and CP wrote and edited the original manuscript. PGP conceived the study and acquired the funds. MG designed the approach, developed, and conceptualized the method, analyzed the data, and wrote the original manuscript. All authors reviewed and approved the manuscript.

